# Regenerating axolotl retinas regrow diverse cell types with modulation by Notch signaling and reconnect to the brain

**DOI:** 10.1101/2022.04.28.489898

**Authors:** Anastasia S. Yandulskaya, Melissa N. Miller, Ronak Ansaripour, Rebecca L. Carrier, James R. Monaghan

## Abstract

Some species successfully repair retinal injuries in contrast to non-regenerative mammalian retina. We show here that the Mexican axolotl salamander regrows its excised retina even in adulthood. During early regeneration, cell proliferation occurred in the retinal pigment epithelium (RPE). All dividing cells expressed *Vimentin*, and some also expressed Müller glia and neural progenitor cell marker *Glast (Slc1a3)*, suggesting that regeneration is driven by RPE-derived retinal progenitor cells. Bulk RNA sequencing showed that genes associated with the extracellular matrix and angiogenesis were upregulated in early-to-mid retinal regeneration. The fully regenerated retina re-established nerve projections to the brain and contained all the original retinal cell types, including Müller glia. Regeneration of cellular diversity may be modulated by Notch signaling, as inhibiting Notch signaling in early regeneration promoted production of rod photoreceptors. Our study highlights the axolotl salamander as an advantageous model of adult tetrapod retinal regeneration and provides insights into its mechanisms.

**Summary:** We demonstrate that adult Mexican axolotl salamanders regenerate retinas after a retinectomy. We also show some cellular and molecular mechanisms that drive axolotl retinal regeneration.

## Introduction

Retinas are susceptible to degeneration that impairs eyesight for life. Humans and other mammals cannot repair this damage, but some vertebrate species regenerate injured retinas. Zebrafish and newt salamanders (Gemberling et al., 2013; Mitashov, 1996) are well-established models of retinal regeneration, but they regrow injured retinas differently. Müller glia repair retinal lesions in zebrafish (Thummel et al., 2008), and retinal pigment epithelium (RPE) regenerates the retina after a retinectomy in Japanese fire-bellied newts (Islam et al., 2014; Thummel et al., 2008). Investigating how other regenerative species regrow their retinas will advance the existing understanding of the mechanisms of retinal regeneration.

The Mexican axolotl (*Ambystoma mexicanum*) presents an intriguing possibility for investigating retinal regeneration. This salamander is a popular model of complex tissue regeneration, most notably of limbs and the spinal cord (Simon & Tanaka, 2013; Tazaki et al., 2017). It can also regenerate portions of ovaries, lungs, the liver, and the brain even in adulthood (Erler et al., 2017; Jensen et al., 2021; Maden et al., 2013; Ohashi et al., 2021). However, the regenerative abilities of axolotl eye tissues at different life stages are either limited or understudied. Axolotl larvae only regenerate lenses for up to two weeks after hatching, after which the ability is lost (Suetsugu-Maki et al., 2012). Juvenile axolotls (4 months) can regenerate removed retinas (Svistunov & Mitashov, 1983), but it remains unknown whether axolotl retinal regeneration persists into adulthood (approximately 1 year of age) or what mechanisms govern it.

We show here that adult paedomorphic axolotls regrow their retinas after a retinectomy, regenerating diverse retinal cell types and re-establishing connections with the brain. Using bulk RNA sequencing, we uncovered genes that are expressed differentially in retinal regeneration. We also show that the Notch signaling pathway may be an important regulator of axolotl retinal regeneration. Our findings establish the axolotl as a model of tetrapod retinal regeneration and provide insights into its molecular mechanisms, which may pave the way towards translating this fascinating ability into treating retinal diseases.

## Methods

### Animal care

Animals were bred in our laboratory or purchased from the Ambystoma Genetic Stock Center (Lexington, KY). The animals were housed as previously described (Farkas, 2015), on a 12:12 light:dark schedule and fed soft salmon pellets. All procedures were approved by Northeastern University Institutional Animal Care and Use Committee. Male and female adult axolotls were used, the term “adult” referring to paedomorphic axolotls who had reached sexual maturity.

### Surgeries and tissue collection

Animals were anesthetized in 0.01% benzocaine prior to all procedures. For eye collection, animals were euthanized in 0.05% benzocaine followed by decapitation.

Cell proliferation: the animals were weighed and injected intraperitoneally with 8.0 ng EdU/g of animal weight in PBS, using 0.33 × 12.7 mm insulin syringes (Exel Int ®). The animals were returned to housing water for 3 hours and then sacrificed for eye collection.

Retinectomies: corneas of the left eyes were punctured with a straight surgical needle and cut halfway around the lens with surgical scissors. The lenses were removed with forceps and the retinas were removed by gently flushing the anterior eye cavities with PBS using a p20 pipette. The animals were returned to housing water in free-standing plastic containers. The right eyes were left uninjured as internal controls. Every retinectomy also involved a lentectomy.

Nerve tracing: we retinectomized 5 adult animals (13-17 cm in total length) and allowed them to regenerate for 367 days. We then anesthetized the animals and cut small V-shaped skin and cartilage flaps in the skulls between the eyes, partially exposing the brain, and labeled the brains with a retrograde nerve tracer cholera toxin subunit B conjugated with Alexa Fluor™ 594 dye (0.5 mg/ml, with approximately 10 µl of final solution per animal). The cartilage flaps were replaced, and the animals were kept under wet lint-free tissues for about 15 minutes with their heads raised to prevent the nerve tracer from leaking out, and then returned to their tanks. The procedure was repeated 3 days later, but the tracer was injected in the existing skull openings without conducting another surgery. The animals were sacrificed at 377 dpi (days post injury), 10 days after the first tracer injection.

DAPT treatment: we soaked small pieces (>0.5mm in diameter) of cotton balls (Thermo Fisher) in 100 mM Notch pathway inhibitor DAPT (Tocris) or its vehicle dimethyl sulfoxide (DMSO; Sigma). We retinectomized the left eyes of 12 male and female adult axolotls (14.5-18 cm in total length). At 15 dpi, we cut small incisions in the corneas of retinectomized eyes and inserted the cotton ball pieces in the intraocular space, as described previously (Nakamura & Chiba, 2007). The treatments were assigned to the animals randomly. The animals were then returned to their housing containers until 101 dpi when they were sacrificed and the left eyes were collected for analysis.

RNA sequencing: the removed (uninjured) retinas were placed in empty 1.5 ml tubes, flash frozen in liquid nitrogen, and stored at -80°C. The animals were returned to their housing containers for 27 days. To collect the regenerating retinas, the animals were sacrificed, the injured eyes were enucleated, and the anterior part of the eyes was carefully trimmed away with surgical scissors. The tissue on the posterior internal side of the exposed eye cups was collected with forceps, placed in 1.5 ml tubes, flash frozen in liquid nitrogen, and stored at -80°C. Both uninjured and regenerating conditions had three biological replicates, with each replicate consisting of four animals, for a total of 12 animals.

### Immunohistochemistry and EdU detection

Collected samples were fixed overnight in 4% paraformaldehyde at 4°C, washed twice for five minutes with phosphate-buffered saline (PBS; Fisher Scientific), and cryoprotected in 30% solution of sucrose in PBS at room temperature until the tissue sank. The samples were then embedded in Optimal Cutting Temperature embedding medium (Fisher Scientific), frozen at -80°C and stored there until use. Tissues were sectioned to 12 µm, washed with deionized water for 5 minutes, washed with PBS for 5 minutes, blocked in 1.5% goat serum for 30 minutes, and incubated in primary antibodies diluted in 1.5% goat serum overnight at 4°C. The sections were then washed with PBS for 5 minutes and incubated in a secondary antibody dilution (1:400 in PBS) for 30 minutes.

Primary antibodies used: rhodopsin (1:400, rabbit, #8710, Cell Signaling Technologies), β-III-tubulin (1:500, mouse, MA1-118, ThermoFisher Scientific). Secondary antibodies used: Alexa Fluor™ 594 goat anti-rabbit IgG (1:400, 1851471, Invitrogen), Alexa Fluor™ 594 goat anti-mouse IgG (1:400, 1851471, Invitrogen).

To detect EdU-positive cells, sections were washed with water and PBS, incubated in Click reaction cocktail (1X tris-buffered saline diluted from stock (Alfa Aesar), 100 mM sodium ascorbate (Acros Organics) in 1X TBS, 4mM CuS04 (Acros Organics) in 1X TBS, and 5 µM AF 594 picolyl azide (Click Chemistry Tools)) for 30 minutes in the dark, washed for five minutes each with PBS, DAPI, PBS, and water, and mounted with 80% glycerol or Slow-fade™ (Invitrogen).

### Version 3 fluorescent in situ hybridization chain reaction (V3 HCR FISH)

We have developed DNA probe sequences for detecting the following cell type markers: *Rhodopsin* (rod photoreceptors) (Osakada et al., 2008), *Rpe65* (the RPE) (Al-Hussaini et al., 2008), *Glast* (also called *Slc1a3;* Müller glia) (Quintero et al., 2016), *Pkcα* (bipolar cells) (Haverkamp et al., 2003), *Lhx1* (horizontal cells) (Poche et al., 2007), and *Glyt1* (amacrine cells) (Nakhai et al., 2007). Probes were also generated for *Vimentin, Notch1, Jag1, Hes1*, and *Hes5*. Probes were designed as described previously (Jensen et al., 2021), using Oligominer (Beliveau et al., 2018) and Bowtie2 (Langmead & Salzberg, 2012) against the 6.0-DD axolotl genome (Nowoshilow et al., 2018; Smith et al., 2019) (http://probegenerator.herokuapp.com/).

Collected samples were embedded in OCT, immediately frozen at -80°C without fixation to limit autofluorescence in the RPE and photoreceptor layers, and stored until use. The samples were sectioned on Superfrost Plus slides at 12 µm thickness on a cryostat (Leica), briefly air-dried at room temperature, and post-fixed with 4% paraformaldehyde for 15 minutes at room temperature. The sections were then washed with PBS (Invitrogen) and dehydrated in 100% ethanol for 15 minutes. Any unused slides were at this point stored in 100% ethanol in sealed LockMailer microscope slides jars at -80°C.

The remaining sections were washed 3×10 minutes with 8% sodium dodecyl sulfate (Fisher Scientific) to further reduce tissue autofluorescence, washed with PBS (Invitrogen), pre-hybridized with pre-warmed hybridization buffer (Molecular Instruments) at 37°C for 15 minutes, and incubated with probes (5 nM in pre-warmed hybridization buffer) at 37°C overnight under parafilm in humidified chambers.

On the next day, the sections were washed with pre-warmed formamide probe wash (Molecular Instruments) 3×15 minutes at 37°C and washed with 5X saline sodium citrate buffer with 0.1% Tween (SSCT) 2×10 minutes at 37°C. Concurrently, hairpins were prepared by aliquoting H1 and H2 hairpins (3 µM; Molecular Instruments) into separate PCR tubes, heating the aliquots at 95°C for 90 seconds in a thermal cycler (Bio-Rad), and cooling them for at least 30 minutes at room temperature in the dark (i.e. in a drawer). The hairpin solution was prepared in the concentration of 1:50, adding 48 µl of amplification buffer (Molecular Instruments) to each pair of cooled 1 µl H1 and H2 hairpins (2 µl of hairpins in total). If several sets of hairpins were used, the volume of the amplification buffer was decreased to maintain the 1:50 dilution ratio. The sections were washed with the amplification buffer for 10 minutes at room temperature and incubated in the hairpin solution under parafilm at room temperature for at least 3 hours or overnight. The sections were then washed with 5X SSCT 3×15 minutes at room temperature, stained with DAPI nuclear stain, washed with PBS and DEPC water, and mounted in Slow-fade™ (Invitrogen) with 24×50-1.5 microscope cover glass (Fisher Scientific).

For multi-round FISH, slides were imaged and then submerged in DEPC water in LockMailer jars to float off coverslips. The sections were treated with DNAse (2000 units/ml) for at least 1 hour at room temperature to wipe away the probes, washed with 60% formamide probe wash for 30 minutes at room temperature, washed with SSCT 2×10 minutes, pre-hybridized, and incubated with new probes overnight at 37°C. The remainder of the staining protocol was followed on the next day.

To preserve RNA in the tissue, RNAse-away spray was used throughout the procedure and all solutions were prepared in deionized diethylpyrocarbonate (DEPC)-treated and autoclaved water. Fresh tissue sections were preferentially used because RNA integrity in sectioned samples can deteriorate during storage. If HCR FISH was combined with EdU detection, the Click reaction was carried out first.

### Bulk RNA sequencing and analysis

RNA isolation and next generation sequencing were performed by Genewiz® (South Plainfield, NJ) with an Illumina® HiSeq® sequencing system. The sequencing parameters were 38 million 150 base pair reads with paired ends. The resulting data was analyzed for differential gene expression and gene ontology.

Differential expression analysis was performed on the Discovery research computing cluster. Raw reads were quality trimmed with Trimmomatic v0.38 (Bolger et al., 2014) and checked with FastQC v0.11.8 (Andrews, 2010). The paired reads were quasi-mapped to the V47 axolotl transcriptome (Nowoshilow & Tanaka, 2020) and quantified at gene level using salmon v0.13.1 (Patro et al., 2017) by the Trinity v2.8.5 (Grabherr et al., 2011) script “align_and_estimate_abundance.pl”. The Trinity package script “abundance_estimates_to_matrix.pl” was used to generate raw counts and TMM-normalized expression matrices. The replicate quality for each group was checked with the Trinity “PtR” script. Differential expression analysis was performed using the “run_DE_analysis.pl” script with the DESeq2 v1.34.0 (Love et al., 2014) method and default parameters. Differential expression results were filtered for padj < 0.001 and log2 fold change > 2 or log2 fold change < -2 to represent genes that were significantly up- or down-regulated in regeneration, respectively. These files were then used in Gene Ontology (GO) analysis to find enriched and depleted GO terms.

GO analysis was performed as follows: Trinotate v3.2.2 (Bryant et al., 2017) was used to generate an annotation report containing top BLAST (Altschul et al., 1990) hits and GO (Ashburner et al., 2000; Gene Ontology, 2021; Mi et al., 2019) assignments from UniProt (UniProt, 2021). GO annotations were analyzed using GOseq v1.46.0 (Young et al., 2010) through the Trinity script “run_GOseq.pl”. The Trinity script “analyze_diff_expr.pl” was run using the GO annotations file to analyze differentially expressed genes for enriched and depleted GO categories in up- and down-regulated genes. The top 25 enriched GO terms in each group were plotted with ggplot2 (Wickham, 2016) in RStudio (RStudio Team, 2020).

### Image quantification and statistical analysis

Images were taken on a Leica DM2500 microscope, a Zeiss LSM 800 confocal microscope, or a Zeiss LSM 880 confocal microscope at Northeastern University Institute for Chemical Imaging of Living Systems (CILS). Confocal images were z-stacks and imaging parameters were the same for all the images within an experiment.

Cell types were quantified in FIJI, using the cell counter plugin. In the 377 dpi experiment, cells of each type were counted within their respective nuclear layer (INL for *Lhx, Pkcα*, and *Glyt1*; ONL for *Rhodopsin*; GCL, INL, and ONL for *Glast*; RPE for *Rpe65*) and normalized to the area of DAPI in those layers. Retinal thickness was measured using the “length” tool, with two measurements per each retinal section.

The data were analyzed with either Student’s two-tailed t-test in Excel or with ANOVA and post-hoc Tukey’s test in JMP 21 statistical software. We used p=0.05 as the significance threshold. Graphs were created using JMP 21. Images were processed in FIJI to improve brightness and contrast, and figures were assembled in Adobe Illustrator 2021.

## Results

### Characterization of the axolotl eye and its retinectomy

We first examined the morphology of the axolotl eye and retina. The axolotl retina follows the conserved retinal structure across vertebrates, containing three nuclear layers and two plexiform layers. The outer and inner nuclear layers in the axolotl retina are very close to each other, so we distinguished them by labeling the outer plexiform layer with a nerve marker β-III-tubulin (BTUBB) (Fig. 1A). Within the eye, the retina can be detected through its expression of rhodopsin (RHO), a gene expressed by rod photoreceptors (Fig. 1B).

**Figure 1:**
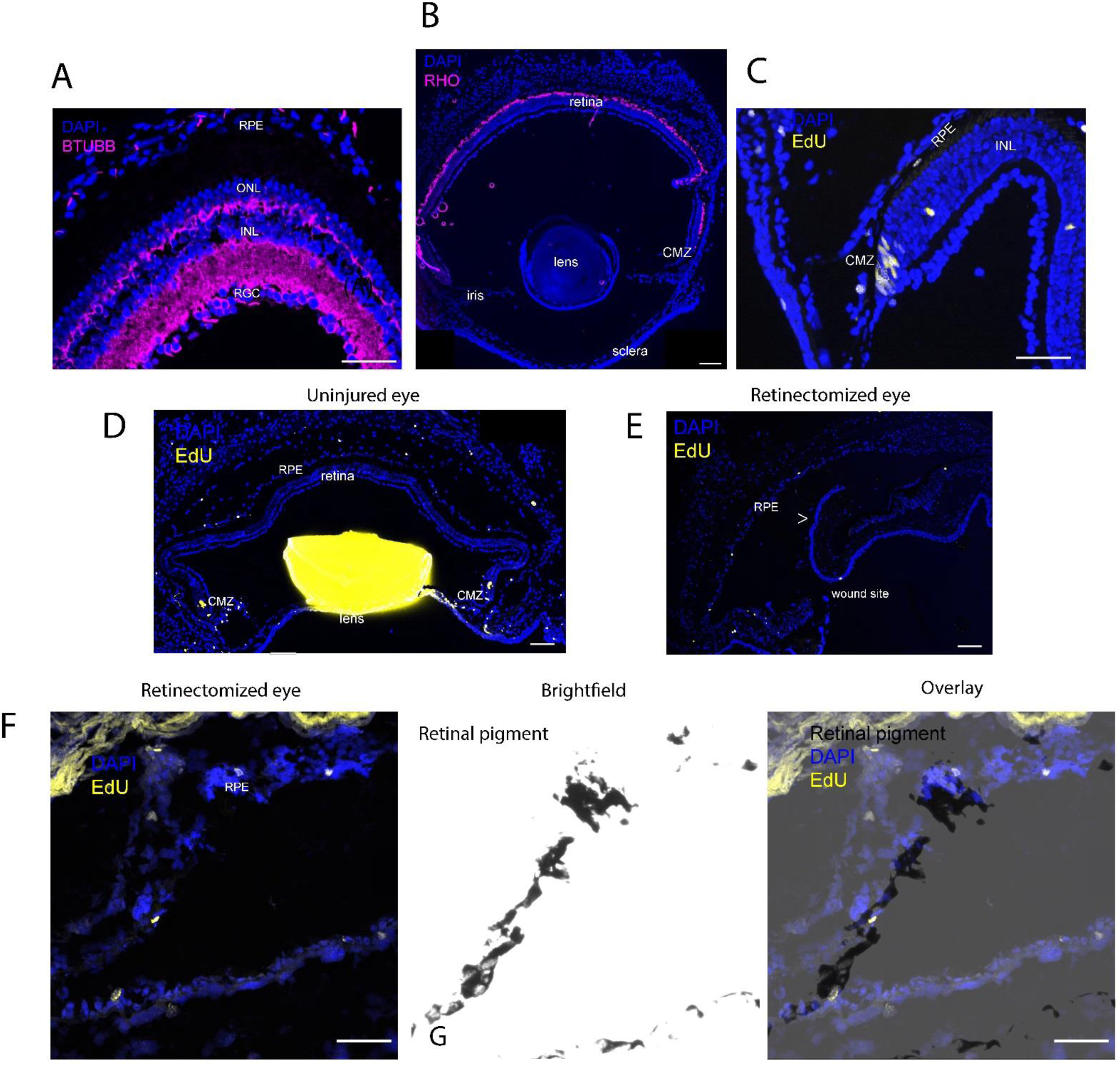
Axolotl eye morphology and retinectomy. (A) The axolotl retina contains three nuclear layers and two plexiform layers that are immunoreactive for the nerve marker BTUBB. The scale bar is 50 μm. (B) A cross-section through the axolotl eye, showing immunostaining for RHO, a photoreceptor marker. (C) EdU+ dividing cells in the homeostatic retina are found in the ciliary marginal zone (CMZ), the retinal pigment epithelium (RPE), and the inner nuclear layer (INL). (D) In an uninjured eye, the retina and the highly autofluorescent lens are present, with cells dividing in the retina and the retinal pigment epithelium (RPE). (E) A retinectomized axolotl eye at 0 dpi. The retinectomy involves a cornea cut, lentectomy, and a retinectomy, preserving the RPE with dividing EdU+ cells. The arrow indicates a scleral flap, which was left in the intravitreal cavity of the immediately collected eye, but it is straightened out in the eyes that are allowed to regenerate. (F) A zoomed-in view of a retinectomized eye. The preserved RPE layer contains EdU+ dividing cells and is discernible in the brightfield view due to its black pigment. The scale bar is 100 μm.

There are three proliferative niches in the homeostatic axolotl retina. Dividing EdU+ cells are found in the ciliary marginal zone (CMZ) and the retinal pigment epithelial layer (RPE), with occasional dividing cells in the inner nuclear layer (Fig. 1C). These three proliferative niches correspond to the three possible pools of retinal progenitor cells: the CMZ, the RPE, and Müller glia. They are affected differently by a retinectomy. Compared to an uninjured eye (Fig. 1D), a retinectomized eye lacks the lens and the retina. The retina, where Müller glia reside, is removed along with the CMZ, although parts of *ora serrata* may remain in the retinal margin (Fig. 1E). The RPE layer may be damaged but is not removed, and it retains dividing cells (Fig. 1E-F). Therefore, the retinectomy is a reliable injury method for the axolotl retina. Retinal regeneration can be assessed using the RHO marker for rod photoreceptors and the BTUBB marker of axons.

### Adult axolotls regenerate their retinas after a retinectomy

We retinectomized the left eyes of 20 adult axolotls and allowed them to regenerate for 15, 35, 49, 65, or 90 days (n=4). We first observed the return of retinal neural layers expressing BTUBB at 49 dpi in one animal out of four (Fig. 2A). By 65 dpi and 90 dpi, the regenerating retinas had reformed the characteristic laminated morphology and distinct nerve layers. Two animals out of four had successfully regenerated their retinas at those time points. The photoreceptor layers (PL), which do not express BTUBB but were visible because of their high autofluorescence, reappeared at 65 dpi. At 90 dpi BTUBB expression was similarly present in the outer plexiform layer (OPL) and inner plexiform layer (IPL) in the regenerated and the uninjured contralateral retinas; the photoreceptor and RPE layers are autofluorescent. The pigment of the RPE layer had also regenerated (Fig. 2B). The lenses never regenerated.

**Figure 2:**
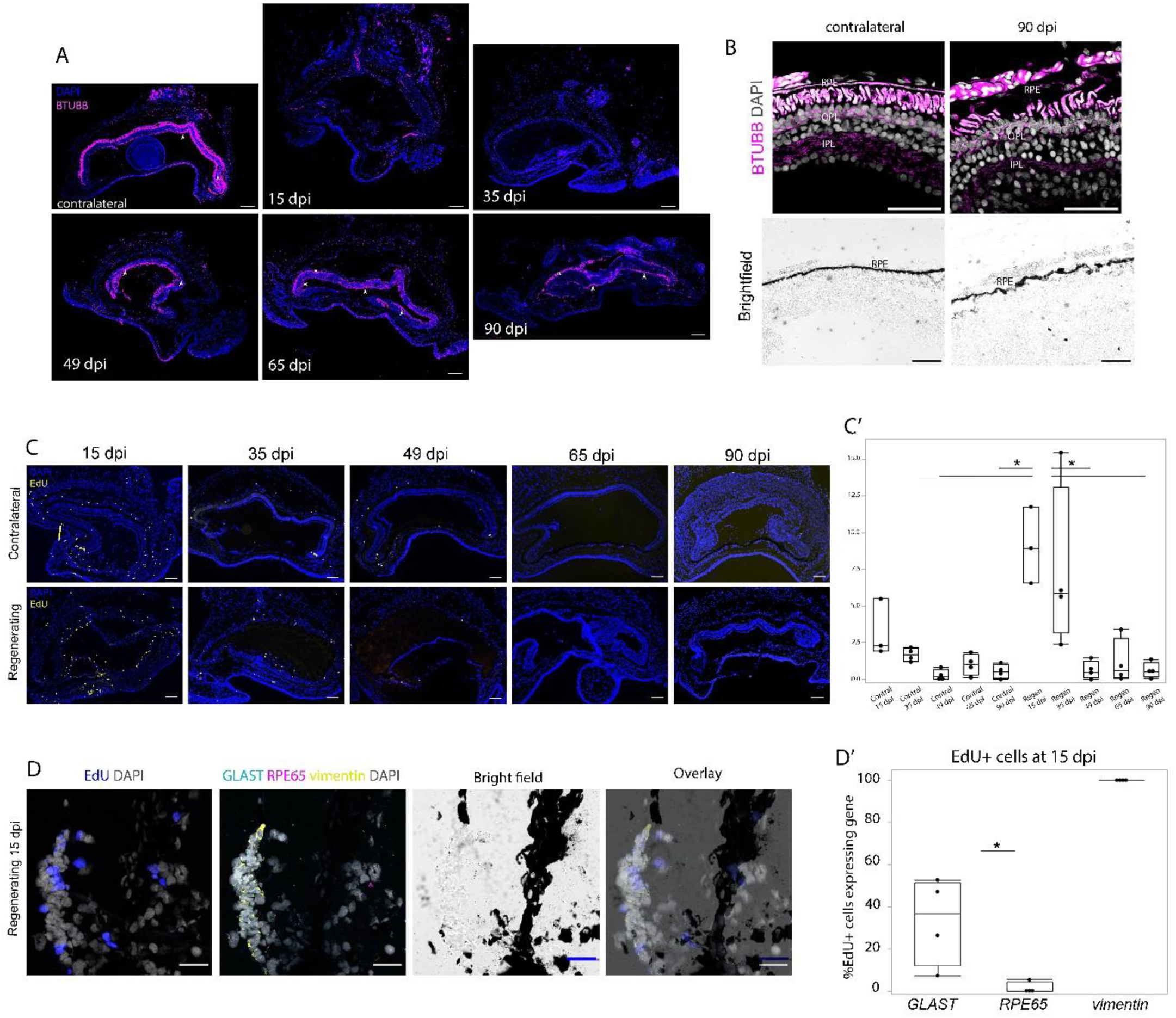
The axolotl retina can regenerate after a retinectomy. (A) Timeline of axolotl retinal regeneration, using immunoreactivity for nerve marker BTUBB to label nerve layers. The retinal structure regenerates within 65 days. (B) The characteristic three-layer retinal lamination with nerve layers (BTUBB) is present both in contralateral and regenerated (90 dpi) retinas in the outer and inner plexiform layers (OPL, IPL). Photoreceptors, the RPE, and the choroid layer are highly autofluorescent. The BTUBB signal is dimmer and DAPI has permeated the photoreceptor and axonal layers because the pictures in (A) and (B) were taken three years apart. (C-C’) EdU+ dividing cells peak at 15 dpi during retinal regeneration, both in the regenerating and contralateral eyes. (D) EdU+ dividing cells express markers of RPE and Müller glia in the regenerating 15 dpi retina, but only of RPE in the contralateral uninjured retina. Arrowheads point to EdU+ cells that express either *Glast* or *Rpe65*. The scale bars are 100 μm in A-C and 50 μm in D.

We then assessed cellular proliferation patterns during regeneration (Fig. 2C). One animal was excluded from the 15 dpi group because its EdU injection had failed. The proportion of EdU+ dividing cells peaked at 15 and 35 dpi. It was not different between these two timepoints (p=0.99). Proliferation in 15 dpi contralateral eyes was not different from contralateral eyes at later time points (p>0.7), but it was not different from either 15 dpi regenerating eyes (p=0.06) or 35 dpi regenerating eyes (p=0.29), which may imply a modest increase in cellular proliferation in the contralateral eyes at 15 dpi. After 35 days, the numbers of dividing cells decreased and were similar in injured and uninjured eyes for the remainder of the regeneration period (Fig. 2C’). The rate of cellular proliferation was highest at 15 dpi and 35 dpi. The regenerating eyes at 15 dpi and 35 dpi had a significantly higher proliferation rate than both contralateral and regenerating eyes at 49 dpi, 65 dpi, and 90 dpi (p<0.01), as well as contralateral eyes at 35 dpi (p=0.02). Proliferation rate was also not different between contralateral and regenerating eyes during later stages of regeneration, at 49 dpi, 65 dpi, and 90 dpi (p>0.99). In regenerating adult axolotl retina, cells divided at the highest rate at 15 dpi and 35 dpi, afterwards decreasing to contralateral uninjured levels. Meanwhile, the contralateral uninjured eye had a small proliferative spike at 15 dpi.

To understand what cell types were dividing at 15 dpi in regenerating eyes, we used HCR FISH to identify EdU+ cells expressing *Glast* (Müller glia marker) and *Rpe65* (RPE marker) in four adult axolotls (Fig. 2D). We also looked for cells positive for *Vim*, which is expressed in Müller glia, neural progenitor cells, and, under some conditions, RPE cells (Guidry et al., 2002). Most dividing EdU+ cells did not express either *Rpe65* or *Glast*, but all expressed *Vim*. 33.3% of dividing cells expressed *Glast*, although they lacked the characteristic elongated morphology of Müller glia. Only 1.3% of dividing cells contained *Rpe65* transcripts, although the presence of black retinal pigment was still substantial in the regenerating retina and co-colocalized with dividing cells (Fig. 2D). *Glast* and *Rpe65* were never co-expressed. Significantly more cells expressed *Glast* than *Rpe65* (p=0.012) (Fig. 2D’). All dividing *Glast+* and *Rpe65+* cells also expressed *Vim*, so we did not examine statistical differences between the expression of *Vim* and the two other genes.

### Regenerated retinas contain diverse cell types and re-establish connections to the brain

We compared the cellular make-up of regenerated and contralateral uninjured axolotl retinas at 377 dpi. We identified the following cell types: rods (*Rho*), bipolar cells (*Lhx1*), horizontal cells (*Pkca*), amacrine cells (*Glyt1*), RPE cells (*Rpe65*), and Müller glia (*Glast)*. We then compared the numbers of those cell types in regenerated and contralateral retinas. The numbers of rods (p=0.44), RPE cells (p=0.13), and bipolar cells (p=0.61) were similar in both uninjured and regenerated retinas (Fig. 3A-C’). However, regenerated retinas contained fewer horizontal (p=0.013) and amacrine cells (p=0.009) and more Müller glia (p=0.013) (Fig. 3D-F’). This experiment suggests that the regenerated axolotl retina regrows the same cell types, some of which are regenerated in different proportions.

**Figure 3:**
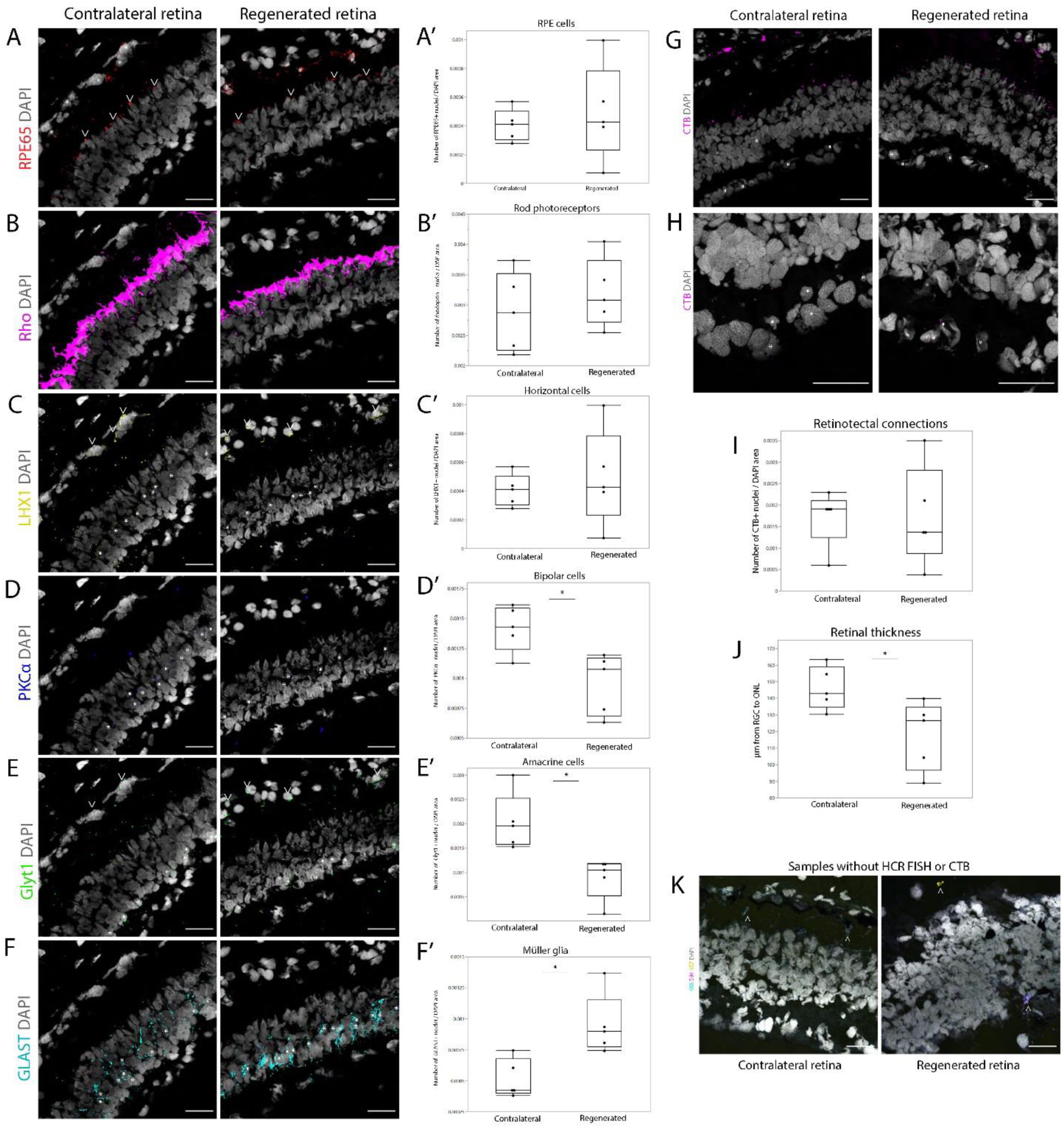
Regenerated retinas (377 dpi) contain diverse cell types and reconnect with the brain. HCR FISH images of cells expressing retinal cell type markers in regenerated and contralateral uninjured retinas, and boxplots showing quantification of those cells: (A-A’) Retinal pigment epithelium (RPE) marker *RPE65*. (B-B’) Rod photoreceptor marker *Rho*. (C-C’) Horizontal cell marker *LHX1*. (D-D’) Bipolar cell marker *Pkc*α. (E-E’) Amacrine cell marker *Glyt1*. (F-F’) Müller glia marker *Glast*. (G-G’) Abundance of RGCs expressing nerve tracer CTB at 20x magnification, at which the data was quantified. The fluorescent signal above the ONL is likely autofluorescence as the sections had not been treated for it. (H-H’) Abundance of RGCs expressing nerve tracer CTB at 40x magnification for better visibility. (I) Boxplot showing retinotectal connectivity of regenerated and contralateral uninjured retinas. (J) Boxplot showing thickness of regenerated and contralateral uninjured neural retinas. (K) Negative control images of V3 HCR FISH, containing only fluorescent hairpins but no probes for detecting gene expression. Asterisks indicate cells expressing a gene, and arrowheads indicate autofluorescence. Scale bars are 50 μm.

We also investigated whether the new retinas had regenerated projections to the brain through the optic nerve. Ten days prior to eye collection, we had injected a retrograde fluorescent nerve tracer cholera toxin subunit B (CTB) that travels from the brain to the retina. All 5 retinectomized eyes had successfully regenerated retinas, and both regenerated and contralateral retinas contained CTB in the RGC layer (Fig. 3G-H). The CTB fluorescence was quantified at 20x magnification (Fig. 3G, I), but it is better visible at 40x magnification (Fig. 3H). The proportion of RGCs containing CTB was similar between contralateral and regenerated retinas (p=0.89) (Fig. 3I), suggesting that regenerated axolotl retinas had successfully restored nerve projections from the retina to the brain. Interestingly, regenerated retinas were thinner than the contralateral ones (p=0.038) (Fig. 3H). Importantly, the HCR FISH protocol quenches the CTB fluorescence (Fig. 3K), and therefore CTB and HCR FISH fluorescence was not present in the retinas at the same time.

### Bulk RNA sequencing of uninjured and regenerating retinas

We conducted bulk RNA sequencing on uninjured and regenerating (27 dpi) retinas, derived from the same 12 adult axolotls, at early-to-mid stages of regeneration when cellular proliferation is still high (Fig. 2C). Both conditions had three biological replicates, each containing tissue pooled from four animals. The pooling was matched between uninjured and regenerating samples (Fig. 4A).

**Figure 4:**
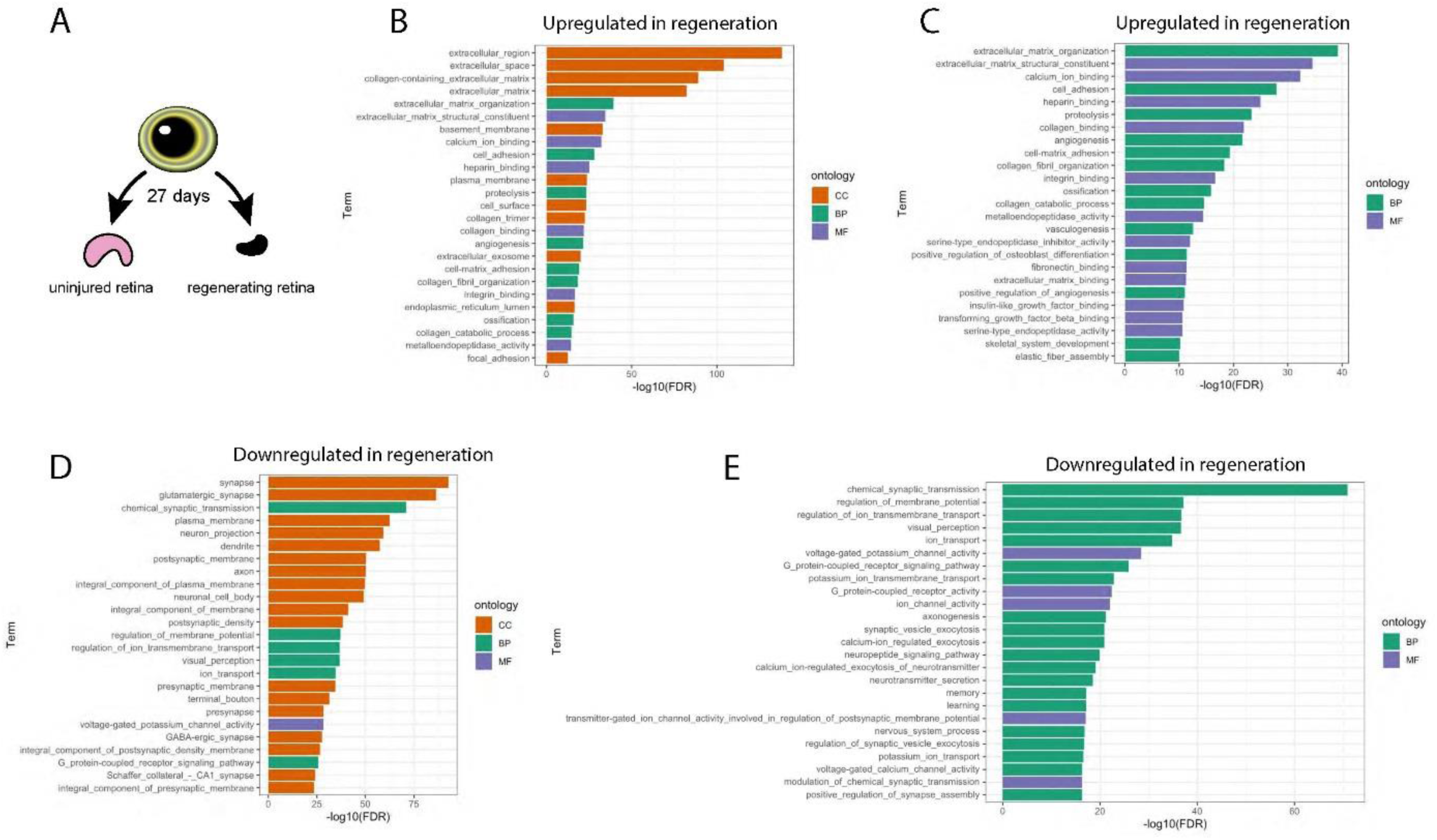
Mechanisms of early-to-mid stage axolotl retinal regeneration. (A) Experimental design of the RNA-Seq analysis. We collected uninjured retinas from 12 adult axolotls and pooled them into 3 samples. We then collected the regenerating tissue from the same retinectomized eyes at 27 dpi, matching the same animals to the 3 samples. (B) Top 25 enriched GO terms that were associated with upregulated genes in retinal regeneration, including “molecular function” (MF), “biological process” (BP), and “cellular component” (CC). (C) Same as in B, excluding “cellular component” category. (D) Top 25 enriched GO terms that were associated with downregulated genes in retinal regeneration, including “molecular function” (MF), “biological process” (BP), and “cellular component” (CC). (E) Same as in D, excluding “cellular component” category.

Genes were identified as differentially expressed if the absolute value of log2fold change was greater than 2. We identified enriched gene ontology (GO) terms in the categories of biological processes (BP), cellular components (CC), and molecular functions (MF). We then identified GO terms that were upregulated and downregulated in regenerating retinal samples. The plots include the top 25 enriched GO terms (Fig. 4).

We identified the GO terms that were associated with genes upregulated in regenerating retinal samples. They were predominantly associated with the extracellular matrix (ECM) (Fig. 4B), such as “extracellular region,” “extracellular matrix,” “collagen-containing extracellular matrix,” “cell-matrix,” “adhesion,” “basement membrane,” “integrin binding,” “and “collagen binding,” highlighting the importance of the extracellular matrix in regeneration.

Most ECM-related GO terms belonged to the “cellular component” category. To ensure that it did not obscure other gene categories, we plotted the top categories for “biological processes” and “molecular processes” only (Fig. 4C), which revealed additional GO terms that were associated with the genes upregulated in regenerating retinal samples. Vascularization of the regenerating retina was represented by GO terms “angiogenesis,” “vasculogenesis,” “positive regulation of angiogenesis,” and “heparin binding.” Signaling pathways were represented by GO terms “insulin-like growth factor binding” (IGF) and “transforming growth factor beta binding” (TGF-β). Interestingly, GO terms “ossification” and “skeletal system development” were also present, even though there are no bones in the retina, possibly implying that bone morphogenesis and retinal tissue growth employ similar molecular mechanisms. GO term “calcium binding” was also present, which may mean that calcium activity regulates regeneration of neural tissue. Finally, this analysis revealed the GO term “fibronectin binding,” which is associated with the extracellular matrix.

We then identified GO terms associated with genes downregulated in regeneration (Fig. 4D). They largely reflected neural activity, such as “synapse,” “regulation of membrane potential,” “chemical synaptic transmission,” and, predictably, “visual perception.” GO terms associated with neuron cell morphology, such as “neuronal cell body,” “axon,” and “neuron projection,” were also present, confirming our previous finding that the retina is not yet regenerated at 27 days, taking more than 35 days (Fig. 2). Plots with “biological processes” and “molecular processes” only (Fig. 4E) revealed GO terms associated with neuronal morphogenesis like “axonogenesis” and “positive regulation of synapse assembly,” suggesting that at 27 dpi, retinal progenitor cells had not yet fully differentiated into neurons.

The differential gene expression analysis of bulk RNA sequencing showed that the extracellular matrix and angiogenesis are important at the early-to-mid stage of axolotl retinal regeneration after a retinectomy. IGF and TGF-β signaling may also regulate retinal regeneration. Neuronal activity and morphogenesis are not restored yet at this stage of regeneration.

### Notch signaling participates in axolotl retinal regeneration

The RNA sequencing analysis revealed differential expression of *Hes1* and *Hes5*, both downstream effectors of the Notch signaling pathway. *Hes1* was upregulated (log2 fold change = 1.76, adjusted p=8.36*10^−45^) and *Hes5* was downregulated (log2 fold change = -2.37, adjusted p=2.37*10^−4^) in the regenerating retina. GO annotation associated *Hes1* with “muscle organ development” (FDR = 0.0030), possibly indicating that axolotl retinal regeneration employs similar mechanisms as organ development. *Hes5* was represented by the following GO terms: “cell adhesion,” “brain development,” “neuron differentiation,” “positive regulation of BMP signaling pathway,” “cartilage development,” “regulation of cell differentiation,” and “negative regulation of oligodendrocyte differentiation” (all FDR values < 0.015). This analysis suggests that during axolotl retinal regeneration, *Hes5* regulates neuronal differentiation.

We qualitatively visualized the expression of *Hes1* and *Hes5*, along two other Notch pathway genes *Notch1* and *Jag1*, in one round and the expression of *Glast, Vim*, and *Rpe65* in the second round of multi-round HCR FISH in a regenerating (28 dpi) eye and the contralateral eye from the same animal (n=1). *Hes1* was present in *Glast*- and *Vim*-expressing Müller glia in the uninjured retina. It was also present in *Glast*-expressing and *Vim*-expressing cells in the regenerating retina that were possibly also expressing *Rpe65* at low levels, although the omnipresence of the *Rpe65* fluorescent stain throughout the tissues complicated definitive identification of *Rpe65*-expressing cells (Fig. 5A, B). Those cells may have been neural progenitor cells or Müller glia that had not yet developed their characteristic elongated morphology. *Hes5* was not detected in the uninjured retina but was present at low levels in the regenerating retina, co-expressed with *Vim* (Fig. 5C, D). *Notch1* and *Jag1*, Notch pathway receptor and ligand, were present throughout the uninjured and regenerating retina. Like in our previous finding (Fig. 2D’), *Vim* was strongly expressed in the regenerating retina. This visualization revealed that *Hes1* is expressed in Müller glia in the uninjured retina and possibly also during regeneration, while *Hes5* is present at low levels in both conditions and its cellular residence remains unclear.

**Figure 5:**
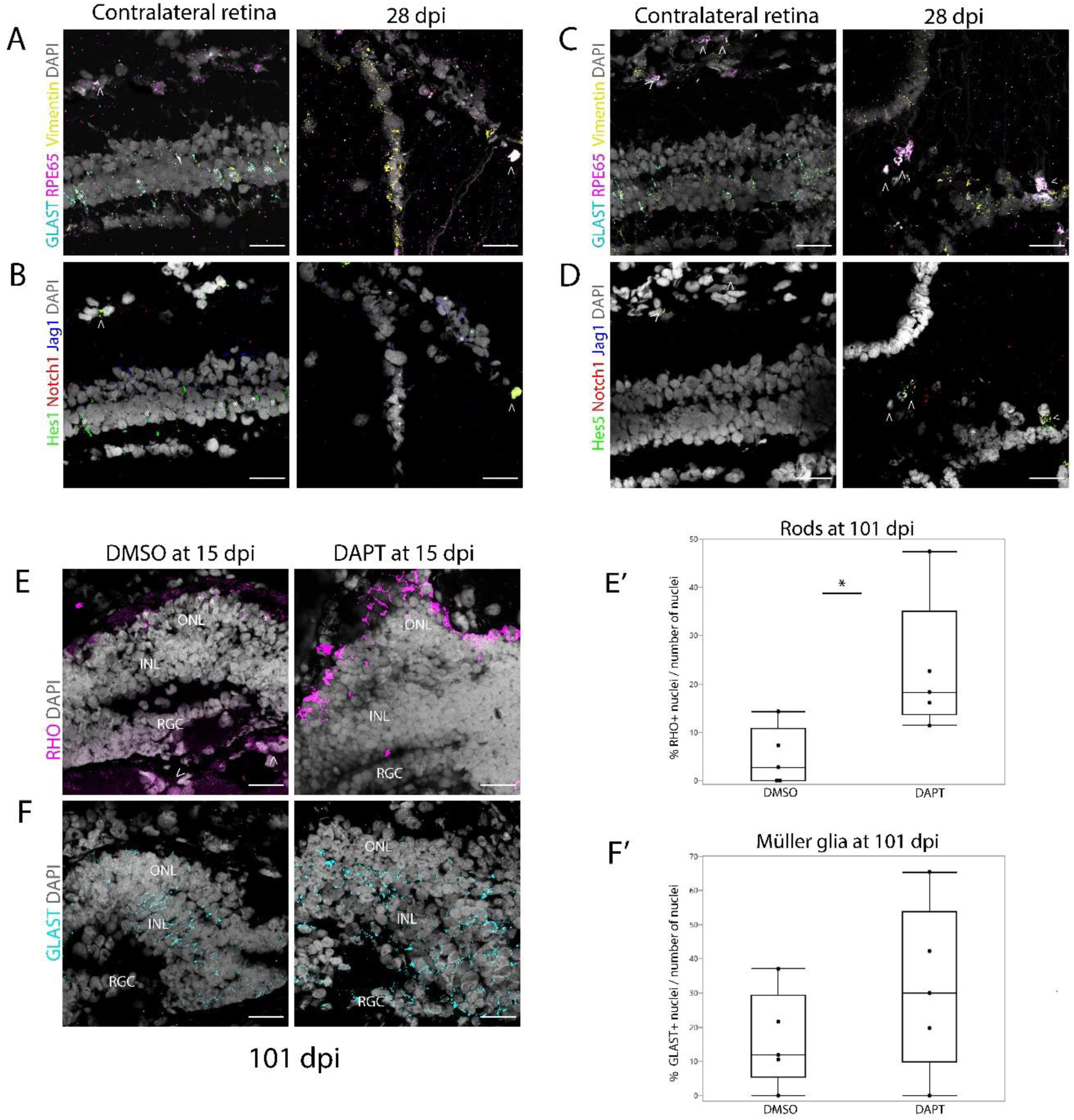
Notch signaling modulates axolotl retinal regeneration. (A, C) Expression of Müller glia marker *Glast*, retinal pigment epithelium (RPE) marker *RPE65*, and *Vim* in a homeostatic retina and regenerating (28 dpi) contralateral retina; (A) was then probed for *Hes1* and (B) for *Hes5*. (B) Expression of *Hes1* and other Notch pathway genes, receptor *Notch1* and ligand *Jag1*, in the same retinal tissue section as in (A). Hes1 is expressed in Müller glia in a homeostatic retina. (D) Expression of *Hes5* and other Notch pathway genes, *Notch1* and *Jag1*, in same tissue sections as (C). (E-E’) Proportion of rod photoreceptors in regenerated (101 dpi) retinas after they were treated with either vehicle or Notch inhibitor DAPT at 15 dpi. (F-F’) Proportion of Müller glia in regenerated (101 dpi) retinas after they were treated with either vehicle or Notch inhibitor DAPT at 15 dpi. Asterisks indicate cells expressing a gene, and arrowheads indicate autofluorescence. Scale bar is 50 μm.

We manipulated Notch signaling during retinal regeneration to better understand its role in neuronal differentiation. We inserted small pieces of cotton balls soaked in either Notch inhibitor DAPT or solvent DMSO into retinectomized eyes (n=4) at 15 dpi, when cellular proliferation peaks (Fig. 2C’). The animals were then allowed to regenerate until 101 dpi. Despite this prolonged regeneration period, regenerated retinal tissue was small and lacked well-defined lamination, possibly due to cotton balls in the eye. One animal was excluded from each treatment group because we could not find sections with regenerated retinas on those samples. Nevertheless, HCR FISH revealed rod marker *Rho* and Müller glia marker *Glast* in the regenerated retinal tissue. DAPT-treated eyes contained more *Rho*+ rod photoreceptors (p=0.041) than solvent-treated eyes (Fig. 5E-E’). The number of *Glast*+ Müller glia was similar in both conditions (p=0.27) (Fig. 5F-F’). These differences in the cellular make-up of regenerated retinal tissue suggest that Notch signaling participates in regulating cell differentiation during axolotl retinal regeneration.

## Discussion

A regenerating retina must produce the original cell types and repair the severed connections with the brain. We show here that adult axolotl salamanders are capable of both, regenerating retinal neural structures within three months after a retinectomy and re-establishing retinotectal projections within a year. To our best knowledge, we have also provided the first visualization of diverse cell types in the axolotl retina, as well as their regeneration.

The axolotl retina contains three possible cellular sources of regeneration: the CMZ, Müller glia, and the RPE. Cellular sources of retinal regeneration vary among species. Zebrafish repair injured retinas using Müller glia (Wan & Goldman, 2016); fire-bellied Japanese newts regenerate missing retinas from the RPE (Mitsuda et al., 2005); post-metamorphic frogs and embryonic chicks have been found to employ all three progenitor cell pools (Langhe et al., 2017; Mitashov & Maliovanova, 1982; Palazzo et al., 2020; Tsonis & Del Rio-Tsonis, 2004; Yoshii et al., 2007). Mammalian retinas do not naturally regenerate, but regenerative responses have been elicited from cultured Müller glia, retinal margin (which anatomically corresponds to the CMZ), and the RPE (Akhtar et al., 2019; Giannelli et al., 2011; Kuwahara et al., 2015; Lawrence et al., 2007; Zhao et al., 1997). Such vast differences in cellular mechanisms of retinal regeneration among species warrant special attention to retinal progenitor cells in the highly regenerative tetrapod axolotl salamander. We observed that all dividing cells in early regeneration expressed *Vimentin* (*Vim*), and one-third also expressed Müller glia marker *Glast. Glast*-positive cells in the regenerating retina lacked the characteristic elongated morphology of Müller glia, but they also did not express the RPE marker *Rpe65* or its black pigment. RPE cells that lose pigment, downregulate RPE65, and express Vimentin are undergoing epithelial-to-mesenchymal transition (EMT) (Tamiya et al., 2010), which marks their new ability to migrate and proliferate (Zhou et al., 2020). This change in the RPE then enables salamander retinal regeneration (Islam et al., 2014). *Glast* may mark dedifferentiating RPE cells, as RPE cells downregulate *Rpe65* in the regenerating newt retina and *Glast* expression has previously been detected in the RPE (Derouiche & Rauen, 1995; Sakami et al., 2005). *Glast* is a Müller glia marker in the homeostatic retina, but it is also expressed in neural progenitor cells in the adult mouse brain (Slezak et al., 2007) and the developing human retina (Walcott & Provis, 2003). We suggest that axolotl retinal regeneration is driven by dedifferentiating cells of the RPE, which then give rise to *Glast*-expressing retinal progenitor cells. Using transgenic reporter axolotl lines in future studies will elucidate the dynamics of RPE reprogramming during axolotl retinal regeneration.

We also found more Müller glia in regenerated retinas than in uninjured ones, which suggests that Müller glia may participate in axolotl retinal regeneration. Alternatively, Müller glia are the last cells born in a developing mammalian retina (Sawant et al., 2019). This pattern may be recapitulated in the regenerating axolotl retina and the numbers of newborn glia may take longer than 377 days to reach baseline levels. However, this is only feasible if axolotl retinal regeneration is driven entirely by the RPE. Developing transgenic reporter axolotls in the future will elucidate the cellular source of retinal regeneration after a retinectomy. Using focal retinal injuries that fully preserve Müller glia populations will further clarify their role in axolotl retinal regeneration.

We observed increased cell proliferation in the intact contralateral eyes. In axolotl regeneration, cells re-enter the cell cycle even at sources distant from the injury. Limb amputation triggers cell-cycle activation in the contralateral, uninjured limbs, but also in other organs (Johnson et al., 2018). After a lung injury, a variety of cell types proliferate in the contralateral, uninjured lung (Jensen et al., 2021). In a regenerating tail, proliferating cells can be detected in the spinal cord as far as 5 mm away from the amputation site (Duerr et al., 2021). It remains unclear what drives this spike in proliferation, but the uninjured contralateral eye may have received growth signals from the brain, which connects the two eyes.

Regenerated axolotl retinas contained all the major retinal cell types and reconnected with the brain. However, they were thinner and did not regenerate the original cell diversity, containing fewer neuronal types of the inner nuclear layer and more Müller glia. This imperfect regeneration of an axolotl neural structure is consistent with a previous observation that the regenerating axolotl brain fails to re-establish pre-existing cytoarchitecture (Amamoto et al., 2016). However, since we compared the regenerated retinas to the uninjured contralateral ones, the regenerated retinas might have been thinner due to a spike in cellular proliferation in contralateral eyes in early regeneration (Fig. 2D’). Finally, retinal regeneration is shaped by neural activity (Chiao et al., 2020). As adult axolotls do not regenerate lenses, re-establishment of correct neuronal networks may suffer from the permanent absence of visual input. Unveiling the molecular mechanisms behind retinal regeneration will help understand how it can be regulated to yield more complete retinal structures.

In species that regenerate the retina from the RPE, this cell monolayer reprograms to give rise to the neural retina using FGF2 (Sakaguchi et al., 1997; Tangeman et al., 2021), and retinal regeneration can be induced by supplying FGF2 to the retinectomized eye (Vergara & Del Rio-Tsonis, 2009). The function of FGF2 may be mediating tissue interactions between the choroid and the RPE, which are important for RPE proliferation (Fronk & Vargis, 2016; Mitsuda et al., 2005). We did not supply any pro-regenerative factors to retinectomized axolotl eyes and saw that the proportion of regenerated retinas varied from 50% (Fig. 2) to 100% (Fig. 3), although foreign bodies in the intraocular space hindered regeneration (Fig. 5). In the future, the success rate of adult axolotl retinal regeneration may be further improved by either carrying out retinectomies with extra care to preserve choroid-RPE interaction, or by experimenting with FGF2 supplementation.

We conducted bulk RNA sequencing of homeostatic and regenerating axolotl retinas. The analysis revealed that at 27 dpi, the regenerating axolotl retina had not yet re-established such essential characteristics as neuronal projections or glutamatergic synaptic signaling. At this timepoint, the injured retina is in early-to-mid regeneration. Genes associated with ECM and angiogenesis were largely upregulated in regenerating retinas. Molecular mechanisms that may regulate early-to-mid stage of axolotl retinal regeneration also include IGF signaling, TGF-β signaling, and calcium binding. IGF signaling participates in retinal progenitor proliferation in fish retina (Becker et al., 2021) and differentiation in mammalian retina (Xia et al., 2018), so it may regulate retinal progenitors in the regenerating axolotl retina. Proteins in the IGF signaling network also regulate angiogenesis (Contois et al., 2012; Delafontaine et al., 2004), possibly linking IGF signaling with angiogenic activity in the regenerating axolotl retina. TGF-β regulates proliferation of retinal progenitor cells in the regenerating zebrafish retina (Sharma et al., 2020), possibly playing a similar role in axolotl retinal regeneration.

The ECM also plays a pivotal role in neural regeneration. Fibronectin is upregulated during early regeneration of newt retina (Ortiz et al., 1992), which agrees with our RNA-Seq data. The ECM of *Xenopus* retinal neuroepithelium cell line promoted axonal outgrowth (Sakaguchi & Radke, 1996), and ECM remodeling in mouse retinal explants improved integration of grafted photoreceptors (Tucker et al., 2008), suggesting that ECM changes may regulate formation of new neuronal networks.

The RNA sequencing analysis also suggested a role for Notch signaling in axolotl retinal regeneration. Notch signaling is a promising candidate for controlling regeneration of retinal cell diversity. The Notch signaling pathway regulates glial and neuronal fate in differentiating neural progenitors, favoring glial fate (Gaiano & Fishell, 2002; Grandbarbe et al., 2003). We used RNA sequencing analysis and visualized its results with HCR FISH to show that two Notch downstream effector genes, *Hes1* and *Hes5*, are differentially expressed during axolotl retinal regeneration. *Hes1* was expressed in Müller glia in an uninjured retina and upregulated in a regenerating retina at 28 dpi. *Hes5* was downregulated during regeneration, although we only detected its weak expression in the retina. These two Notch pathway genes have distinct functions during retinogenesis: *Hes1* maintains retinal progenitor cells in an undifferentiated state, while *Hes5* promotes their glial fate and inhibits their differentiation into neurons (Hojo et al., 2000). *Hes5*, but not *Hes1*, was upregulated in the embryonic chick retina after an excitotoxic injury (Hayes et al., 2007). Our analysis shows the opposite pattern, although it may have been carried out at a different stage of retinal regeneration and employed a different injury method (3 days post excitotoxic injury in chicks vs. 27 days post retinectomy in axolotls). Upregulation of *Hes1* may indicate that it contributes to maintaining a pool of progenitor cells that are necessary for regrowing the retina. Downregulation of *Hes5* suggests that at 28 dpi, retinal progenitor cells may be committing to a specific fate, emphasizing the role of Notch signaling in retinal differentiation.

Notch signaling favors glial and progenitor cell fate in the retina (Jadhav et al., 2006). Inhibiting Notch signaling promotes regeneration of neurons, such as hair cells and motor neurons of the spinal cord (Dias et al., 2012; Mizutari et al., 2013). In the retina, inhibition of Notch triggers a regenerative response (Conner et al., 2014; Elsaeidi et al., 2018; Hayes et al., 2007), and its dynamic regulation is required for regeneration of the zebrafish retina (Campbell et al., 2022). In Japanese fire-bellied newts, intraocular inhibition of Notch accelerated formation of neurons during early regeneration of the retina (Nakamura & Chiba, 2007). We show here that early inhibition of Notch signaling also impacts retinal regeneration in the long term, driving production of rod photoreceptors. If Müller glia self-renew in axolotls like in zebrafish (Langhe & Pearson, 2020; Meyers et al., 2012), they may have compensated Notch inhibition-induced deficit in their numbers during regeneration. Our finding emphasizes Notch signaling as an important regulator of cell diversity during retinal regeneration.

The axolotl salamander is a useful model of retinal regeneration because it regenerates its retina even in adulthood and accommodates techniques of assessing and manipulating gene expression. Much larger than zebrafish, axolotls are amenable to retinectomies and more focal injuries like light or excitotoxicity; studying retinal responses to different injuries may help clarify how different progenitor cells function in regeneration. Unlike some newt species, axolotls are also easy to breed in a laboratory and to develop into transgenic lines (Joven et al., 2019), which will aid future studies of their retinal regeneration. In addition, retinectomies - often-used models of retinal injury in salamanders - require removing both the retina and the lens. Newts can regenerate their lenses, but axolotls lose this ability soon after hatching (Suetsugu-Maki et al., 2012). Therefore, retinectomized axolotl eyes only regrow the retina without concurrently regenerating the lens, and any regenerative processes, such as cell proliferation, can likely be attributed to retinal regeneration. Studies on behavior and gene expression of axolotl retinal progenitors will enrich our understanding of retinal regeneration across the phylogeny, harness the regenerative power of those cell types in the human retina, and, in the future, help place humans among species that can repair their retinas.

## Acknowledgements

The funding was provided by the Retina Research Foundation (to JRM) and NSF award #1606128 (to RLC and JRM). The Ambystoma Genetic Stock Center (Lexington, KY) is supported by the NIH grant P40-OD019794. The funding sources were not involved in the study design or execution.

The high-performance computing resources for RNA-Seq analysis were provided by the Northeastern Discovery cluster at the Massachusetts Green-High Performance Computing Center in Holyoke, MA. We thank the Institute for Chemical Imaging of Living Systems at Northeastern University for consultation and imaging support. We also thank Dr. Alexander Lovely for assistance with imaging and Jackson Griffiths for insightful comments on the manuscript.

## Abbreviations

RPE: retinal pigment epithelium
ONL: outer nuclear layer
INL: inner nuclear layer
RGC: retinal ganglion cells
OPL: outer plexiform layer
IPL: inner plexiform layer
CTB: cholera toxin subunit B
GFP: green fluorescent protein
ECM: the extracellular matrix
IGF: insulin-like growth factor
TGF-β: transforming growth factor beta

## References

Akhtar, T., Xie, H., Khan, M. I., Zhao, H., Bao, J., Zhang, M., & Xue, T. (2019). Accelerated photoreceptor differentiation of hiPSC-derived retinal organoids by contact co-culture with retinal pigment epithelium. Stem Cell Res, 39, 101491. https://doi.org/10.1016/j.scr.2019.101491

Al-Hussaini, H., Kam, J. H., Vugler, A., Semo, M., & Jeffery, G. (2008). Mature retinal pigment epithelium cells are retained in the cell cycle and proliferate in vivo. Mol Vis, 14, 1784–1791. https://www.ncbi.nlm.nih.gov/pubmed/18843376

Altschul, S. F., Gish, W., Miller, W., Myers, E. W., & Lipman, D. J. (1990). Basic local alignment search tool. J Mol Biol, 215(3), 403–410. https://doi.org/10.1016/S0022-2836(05)80360-2

Amamoto, R., Huerta, V. G., Takahashi, E., Dai, G., Grant, A. K., Fu, Z., & Arlotta, P. (2016). Adult axolotls can regenerate original neuronal diversity in response to brain injury. Elife, 5. https://doi.org/10.7554/eLife.13998

Andrews, S. (2010). FastQC: a quality control tool for high throughput sequence data.

Ashburner, M., Ball, C. A., Blake, J. A., Botstein, D., Butler, H., Cherry, J. M., Davis, A. P., Dolinski, K., Dwight, S. S., Eppig, J. T., Harris, M. A., Hill, D. P., Issel-Tarver, L., Kasarskis, A., Lewis, S., Matese, J. C., Richardson, J. E., Ringwald, M., Rubin, G. M., … Consortium, G. O. (2000). Gene Ontology: tool for the unification of biology. Nature Genetics, 25(1), 25–29. https://doi.org/Doi10.1038/75556

Becker, C., Lust, K., & Wittbrodt, J. (2021). Igf signaling couples retina growth with body growth by modulating progenitor cell division. Development, 148(7). https://doi.org/10.1242/dev.199133

Beliveau, B. J., Kishi, J. Y., Nir, G., Sasaki, H. M., Saka, S. K., Nguyen, S. C., Wu, C. T., & Yin, P. (2018). OligoMiner provides a rapid, flexible environment for the design of genome-scale oligonucleotide in situ hybridization probes. Proc Natl Acad Sci U S A, 115(10), E2183–E2192. https://doi.org/10.1073/pnas.1714530115

Bolger, A. M., Lohse, M., & Usadel, B. (2014). Trimmomatic: a flexible trimmer for Illumina sequence data. Bioinformatics, 30(15), 2114–2120. https://doi.org/10.1093/bioinformatics/btu170

Bryant, D. M., Johnson, K., DiTommaso, T., Tickle, T., Couger, M. B., Payzin-Dogru, D., Lee, T. J., Leigh, N. D., Kuo, T. H., Davis, F. G., Bateman, J., Bryant, S., Guzikowski, A. R., Tsai, S. L., Coyne, S., Ye, W. W., Freeman, R. M., Jr., Peshkin, L., Tabin, C. J., Whited, J. L. (2017). A Tissue-Mapped Axolotl De Novo Transcriptome Enables Identification of Limb Regeneration Factors. Cell Rep, 18(3), 762–776. https://doi.org/10.1016/j.celrep.2016.12.063

Campbell, L. J., Levendusky, J. L., Steines, S. A., & Hyde, D. R. (2022). Retinal regeneration requires dynamic Notch signaling. Neural Regen Res, 17(6), 1199–1209. https://doi.org/10.4103/1673-5374.327326

Chiao, C. C., Lin, C. I., & Lee, M. J. (2020). Multiple Approaches for Enhancing Neural Activity to Promote Neurite Outgrowth of Retinal Explants. Methods Mol Biol, 2092, 65–75. https://doi.org/10.1007/978-1-0716-0175-4_6

Conner, C., Ackerman, K. M., Lahne, M., Hobgood, J. S., & Hyde, D. R. (2014). Repressing notch signaling and expressing TNFalpha are sufficient to mimic retinal regeneration by inducing Muller glial proliferation to generate committed progenitor cells. J Neurosci, 34(43), 14403–14419. https://doi.org/10.1523/JNEUROSCI.0498-14.2014

Contois, L. W., Nugent, D. P., Caron, J. M., Cretu, A., Tweedie, E., Akalu, A., Liebes, L., Friesel, R., Rosen, C., Vary, C., & Brooks, P. C. (2012). Insulin-like growth factor binding protein-4 differentially inhibits growth factor-induced angiogenesis. J Biol Chem, 287(3), 1779–1789. https://doi.org/10.1074/jbc.M111.267732

Delafontaine, P., Song, Y. H., & Li, Y. (2004). Expression, regulation, and function of IGF-1, IGF-1R, and IGF-1 binding proteins in blood vessels. Arterioscler Thromb Vasc Biol, 24(3), 435–444. https://doi.org/10.1161/01.ATV.0000105902.89459.09

Derouiche, A., & Rauen, T. (1995). Coincidence of L-glutamate/L-aspartate transporter (GLAST) and glutamine synthetase (GS) immunoreactions in retinal glia: evidence for coupling of GLAST and GS in transmitter clearance. J Neurosci Res, 42(1), 131–143. https://doi.org/10.1002/jnr.490420115

Dias, T. B., Yang, Y. J., Ogai, K., Becker, T., & Becker, C. G. (2012). Notch signaling controls generation of motor neurons in the lesioned spinal cord of adult zebrafish. J Neurosci, 32(9), 3245–3252. https://doi.org/10.1523/JNEUROSCI.6398-11.2012

Duerr, T. J., Jeon, E. K., Wells, K. M., Villanueva, A., Seifert, A. W., McCusker, C. D., & Monaghan, J. R. (2021). A constitutively expressed fluorescence ubiquitin cell cycle indicator (FUCCI) in axolotls for studying tissue regeneration. bioRxiv, 2021.2003.2030.437716. https://doi.org/10.1101/2021.03.30.437716

Elsaeidi, F., Macpherson, P., Mills, E. A., Jui, J., Flannery, J. G., & Goldman, D. (2018). Notch Suppression Collaborates with Ascl1 and Lin28 to Unleash a Regenerative Response in Fish Retina, But Not in Mice. J Neurosci, 38(9), 2246–2261. https://doi.org/10.1523/JNEUROSCI.2126-17.2018

Erler, P., Sweeney, A., & Monaghan, J. R. (2017). Regulation of Injury-Induced Ovarian Regeneration by Activation of Oogonial Stem Cells. Stem Cells, 35(1), 236–247. https://doi.org/10.1002/stem.2504

Fronk, A. H., & Vargis, E. (2016). Methods for culturing retinal pigment epithelial cells: a review of current protocols and future recommendations. J Tissue Eng, 7, 2041731416650838. https://doi.org/10.1177/2041731416650838

Gaiano, N., & Fishell, G. (2002). The role of notch in promoting glial and neural stem cell fates. Annu Rev Neurosci, 25, 471–490. https://doi.org/10.1146/annurev.neuro.25.030702.130823

Gemberling, M., Bailey, T. J., Hyde, D. R., & Poss, K. D. (2013). The zebrafish as a model for complex tissue regeneration. Trends in Genetics, 29(11), 611–620. https://doi.org/10.1016/j.tig.2013.07.003

Gene Ontology, C. (2021). The Gene Ontology resource: enriching a GOld mine. Nucleic Acids Res, 49(D1), D325–D334. https://doi.org/10.1093/nar/gkaa1113

Giannelli, S. G., Demontis, G. C., Pertile, G., Rama, P., & Broccoli, V. (2011). Adult human Muller glia cells are a highly efficient source of rod photoreceptors. Stem Cells, 29(2), 344–356. https://doi.org/10.1002/stem.579

Grabherr, M. G., Haas, B. J., Yassour, M., Levin, J. Z., Thompson, D. A., Amit, I., Adiconis, X., Fan, L., Raychowdhury, R., Zeng, Q., Chen, Z., Mauceli, E., Hacohen, N., Gnirke, A., Rhind, N., di Palma, F., Birren, B. W., Nusbaum, C., Lindblad-Toh, K., … Regev, A. (2011). Full-length transcriptome assembly from RNA-Seq data without a reference genome. Nat Biotechnol, 29(7), 644–652. https://doi.org/10.1038/nbt.1883

Grandbarbe, L., Bouissac, J., Rand, M., Hrabe de Angelis, M., Artavanis-Tsakonas, S., & Mohier, E. (2003). Delta-Notch signaling controls the generation of neurons/glia from neural stem cells in a stepwise process. Development, 130(7), 1391–1402. https://doi.org/10.1242/dev.00374

Guidry, C., Medeiros, N. E., & Curcio, C. A. (2002). Phenotypic variation of retinal pigment epithelium in age-related macular degeneration. Invest Ophthalmol Vis Sci, 43(1), 267–273. https://www.ncbi.nlm.nih.gov/pubmed/11773041

Wickham, H. (2016). ggplot2: Elegant Graphics for Data Analysis. Springer-Verlag New York.

Haverkamp, S., Haeseleer, F., & Hendrickson, A. (2003). A comparison of immunocytochemical markers to identify bipolar cell types in human and monkey retina. Vis Neurosci, 20(6), 589–600. https://doi.org/10.1017/s0952523803206015

Hayes, S., Nelson, B. R., Buckingham, B., & Reh, T. A. (2007). Notch signaling regulates regeneration in the avian retina. Dev Biol, 312(1), 300–311. https://doi.org/10.1016/j.ydbio.2007.09.046

Hojo, M., Ohtsuka, T., Hashimoto, N., Gradwohl, G., Guillemot, F., & Kageyama, R. (2000). Glial cell fate specification modulated by the bHLH gene Hes5 in mouse retina. Development, 127(12), 2515–2522. https://www.ncbi.nlm.nih.gov/pubmed/10821751

Islam, M. R., Nakamura, K., Casco-Robles, M. M., Kunahong, A., Inami, W., Toyama, F., Maruo, F., & Chiba, C. (2014). The newt reprograms mature RPE cells into a unique multipotent state for retinal regeneration. Sci Rep, 4, 6043. https://doi.org/10.1038/srep06043

Jadhav, A. P., Cho, S. H., & Cepko, C. L. (2006). Notch activity permits retinal cells to progress through multiple progenitor states and acquire a stem cell property. Proc Natl Acad Sci U S A, 103(50), 18998–19003. https://doi.org/10.1073/pnas.0608155103

Jensen, T. B., Giunta, P., Schultz, N. G., Griffiths, J. M., Duerr, T. J., Kyeremateng, Y., Wong, H., Adesina, A., & Monaghan, J. R. (2021). Lung injury in axolotl salamanders induces an organ-wide proliferation response. Dev Dyn, 250(6), 866–879. https://doi.org/10.1002/dvdy.315

Johnson, K., Bateman, J., DiTommaso, T., Wong, A. Y., & Whited, J. L. (2018). Systemic cell cycle activation is induced following complex tissue injury in axolotl. Dev Biol, 433(2), 461–472. https://doi.org/10.1016/j.ydbio.2017.07.010

Joven, A., Elewa, A., & Simon, A. (2019). Model systems for regeneration: salamanders. Development, 146(14). https://doi.org/10.1242/dev.167700

Kuwahara, A., Ozone, C., Nakano, T., Saito, K., Eiraku, M., & Sasai, Y. (2015). Generation of a ciliary margin-like stem cell niche from self-organizing human retinal tissue. Nat Commun, 6, 6286. https://doi.org/10.1038/ncomms7286

Langhe, R., Chesneau, A., Colozza, G., Hidalgo, M., Ail, D., Locker, M., & Perron, M. (2017). Muller glial cell reactivation in Xenopus models of retinal degeneration. Glia, 65(8), 1333–1349. https://doi.org/10.1002/glia.23165

Langhe, R., & Pearson, R. A. (2020). Rebuilding the Retina: Prospects for Muller Glial-mediated Self-repair. Curr Eye Res, 45(3), 349–360. https://doi.org/10.1080/02713683.2019.1669665

Langmead, B., & Salzberg, S. L. (2012). Fast gapped-read alignment with Bowtie 2. Nat Methods, 9(4), 357–359. https://doi.org/10.1038/nmeth.1923

Lawrence, J. M., Singhal, S., Bhatia, B., Keegan, D. J., Reh, T. A., Luthert, P. J., Khaw, P. T., & Limb, G. A. (2007). MIO-M1 cells and similar Muller glial cell lines derived from adult human retina exhibit neural stem cell characteristics. Stem Cells, 25(8), 2033–2043. https://doi.org/10.1634/stemcells.2006-0724

Love, M. I., Huber, W., & Anders, S. (2014). Moderated estimation of fold change and dispersion for RNA-seq data with DESeq2. Genome Biol, 15(12), 550. https://doi.org/10.1186/s13059-014-0550-8

Maden, M., Manwell, L. A., & Ormerod, B. K. (2013). Proliferation zones in the axolotl brain and regeneration of the telencephalon. Neural Dev, 8, 1. https://doi.org/10.1186/1749-8104-8-1

Meyers, J. R., Hu, L., Moses, A., Kaboli, K., Papandrea, A., & Raymond, P. A. (2012). beta-catenin/Wnt signaling controls progenitor fate in the developing and regenerating zebrafish retina. Neural Dev, 7, 30. https://doi.org/10.1186/1749-8104-7-30

Mi, H., Muruganujan, A., Ebert, D., Huang, X., & Thomas, P. D. (2019). PANTHER version 14: more genomes, a new PANTHER GO-slim and improvements in enrichment analysis tools. Nucleic Acids Res, 47(D1), D419–D426. https://doi.org/10.1093/nar/gky1038

Mitashov, V. I. (1996). Mechanisms of retina regeneration in urodeles. International Journal of Developmental Biology, 40(4), 833–844. <Go to ISI>://WOS:A1996VE44100028

Mitashov, V. I., & Maliovanova, S. D. (1982). [Cellular proliferative potentials of the pigment and ciliated epithelium of the eye in clawed toads normally and during regeneration]. Ontogenez, 13(3), 228–234. https://www.ncbi.nlm.nih.gov/pubmed/7099515 (Proliferativnye potentsii kletok pigmentnogo i tsiliarnogo epiteliaa glaza shportsevykh liagushek v norme i pri regeneratsii.)

Mitsuda, S., Yoshii, C., Ikegami, Y., & Araki, M. (2005). Tissue interaction between the retinal pigment epithelium and the choroid triggers retinal regeneration of the newt Cynops pyrrhogaster. Dev Biol, 280(1), 122–132. https://doi.org/10.1016/j.ydbio.2005.01.009

Mizutari, K., Fujioka, M., Hosoya, M., Bramhall, N., Okano, H. J., Okano, H., & Edge, A. S. (2013). Notch inhibition induces cochlear hair cell regeneration and recovery of hearing after acoustic trauma. Neuron, 77(1), 58–69. https://doi.org/10.1016/j.neuron.2012.10.032

Nakamura, K., & Chiba, C. (2007). Evidence for Notch signaling involvement in retinal regeneration of adult newt. Brain Res, 1136(1), 28–42. https://doi.org/10.1016/j.brainres.2006.12.032

Nakhai, H., Sel, S., Favor, J., Mendoza-Torres, L., Paulsen, F., Duncker, G. I., & Schmid, R. M. (2007). Ptf1a is essential for the differentiation of GABAergic and glycinergic amacrine cells and horizontal cells in the mouse retina. Development, 134(6), 1151–1160. https://doi.org/10.1242/dev.02781

Nowoshilow, S., Schloissnig, S., Fei, J. F., Dahl, A., Pang, A. W. C., Pippel, M., Winkler, S., Hastie, A. R., Young, G., Roscito, J. G., Falcon, F., Knapp, D., Powell, S., Cruz, A., Cao, H., Habermann, B., Hiller, M., Tanaka, E. M., & Myers, E. W. (2018). The axolotl genome and the evolution of key tissue formation regulators. Nature, 554(7690), 50–55. https://doi.org/10.1038/nature25458

Nowoshilow, S., & Tanaka, E. M. (2020). Introducing www.axolotl-omics.org -an integrated -omics data portal for the axolotl research community. Exp Cell Res, 394(1), 112143. https://doi.org/10.1016/j.yexcr.2020.112143

Ohashi, A., Saito, N., Kashimoto, R., Furukawa, S., Yamamoto, S., & Satoh, A. (2021). Axolotl liver regeneration is accomplished via compensatory congestion mechanisms regulated by ERK signaling after partial hepatectomy. Dev Dyn, 250(6), 838–851. https://doi.org/10.1002/dvdy.262

Ortiz, J. R., Vigny, M., Courtois, Y., & Jeanny, J. C. (1992). Immunocytochemical study of extracellular matrix components during lens and neural retina regeneration in the adult newt. Exp Eye Res, 54(6), 861–870. https://doi.org/10.1016/0014-4835(92)90149-m

Osakada, F., Ikeda, H., Mandai, M., Wataya, T., Watanabe, K., Yoshimura, N., Akaike, A., Sasai, Y., & Takahashi, M. (2008). Toward the generation of rod and cone photoreceptors from mouse, monkey and human embryonic stem cells. Nat Biotechnol, 26(2), 215–224. https://doi.org/10.1038/nbt1384

Palazzo, I., Deistler, K., Hoang, T. V., Blackshaw, S., & Fischer, A. J. (2020). NF-kappaB signaling regulates the formation of proliferating Muller glia-derived progenitor cells in the avian retina. Development, 147(10). https://doi.org/10.1242/dev.183418

Patro, R., Duggal, G., Love, M. I., Irizarry, R. A., & Kingsford, C. (2017). Salmon provides fast and bias-aware quantification of transcript expression. Nat Methods, 14(4), 417–419. https://doi.org/10.1038/nmeth.4197

Poche, R. A., Kwan, K. M., Raven, M. A., Furuta, Y., Reese, B. E., & Behringer, R. R. (2007). Lim1 is essential for the correct laminar positioning of retinal horizontal cells. J Neurosci, 27(51), 14099–14107. https://doi.org/10.1523/JNEUROSCI.4046-07.2007

Quintero, H., Gomez-Montalvo, A. I., & Lamas, M. (2016). MicroRNA changes through Muller glia dedifferentiation and early/late rod photoreceptor differentiation. Neuroscience, 316, 109–121. https://doi.org/10.1016/j.neuroscience.2015.12.025

RStudio Team (2020). RStudio: Integrated Development for R. RStudio, PBC, Boston, MA http://www.rstudio.com/

Sakaguchi, D. S., Janick, L. M., & Reh, T. A. (1997). Basic fibroblast growth factor (FGF-2) induced transdifferentiation of retinal pigment epithelium: generation of retinal neurons and glia. Dev Dyn, 209(4), 387–398. https://doi.org/10.1002/(SICI)1097-0177(199708)209:4<387::AID-AJA6>3.0.CO;2-E

Sakaguchi, D. S., & Radke, K. (1996). Beta 1 integrins regulate axon outgrowth and glial cell spreading on a glial-derived extracellular matrix during development and regeneration. Brain Res Dev Brain Res, 97(2), 235–250. https://doi.org/10.1016/s0165-3806(96)00142-3

Sakami, S., Hisatomi, O., Sakakibara, S., Liu, J., Reh, T. A., & Tokunaga, F. (2005). Downregulation of Otx2 in the dedifferentiated RPE cells of regenerating newt retina. Brain Res Dev Brain Res, 155(1), 49–59. https://doi.org/10.1016/j.devbrainres.2004.11.008

Sawant, O. B., Jidigam, V. K., Fuller, R. D., Zucaro, O. F., Kpegba, C., Yu, M., Peachey, N. S., & Rao, S. (2019). The circadian clock gene Bmal1 is required to control the timing of retinal neurogenesis and lamination of Muller glia in the mouse retina. FASEB J, 33(8), 8745–8758. https://doi.org/10.1096/fj.201801832RR

Sharma, P., Gupta, S., Chaudhary, M., Mitra, S., Chawla, B., Khursheed, M. A., Saran, N. K., & Ramachandran, R. (2020). Biphasic Role of Tgf-beta Signaling during Muller Glia Reprogramming and Retinal Regeneration in Zebrafish. iScience, 23(2), 100817. https://doi.org/10.1016/j.isci.2019.100817

Simon, A., & Tanaka, E. M. (2013). Limb regeneration. Wiley Interdiscip Rev Dev Biol, 2(2), 291–300. https://doi.org/10.1002/wdev.73

Slezak, M., Goritz, C., Niemiec, A., Frisen, J., Chambon, P., Metzger, D., & Pfrieger, F. W. (2007). Transgenic mice for conditional gene manipulation in astroglial cells. Glia, 55(15), 1565–1576. https://doi.org/10.1002/glia.20570

Smith, J. J., Timoshevskaya, N., Timoshevskiy, V. A., Keinath, M. C., Hardy, D., & Voss, S. R. (2019). A chromosome-scale assembly of the axolotl genome. Genome Res, 29(2), 317–324. https://doi.org/10.1101/gr.241901.118

Suetsugu-Maki, R., Maki, N., Nakamura, K., Sumanas, S., Zhu, J., Del Rio-Tsonis, K., & Tsonis, P. A. (2012). Lens regeneration in axolotl: new evidence of developmental plasticity. BMC Biol, 10, 103. https://doi.org/10.1186/1741-7007-10-103

Svistunov, S. A., & Mitashov, V. I. (1983). [Proliferative activity of the pigment epithelium and regenerating retinal cells in Ambystoma mexicanum]. Ontogenez, 14(6), 597–606. https://www.ncbi.nlm.nih.gov/pubmed/6657169 (Proliferativnaia aktivnost’ kletok pigmentnogo epiteliia i regeneriruiushchei setchatki u Ambystoma mexicanum.)

Tamiya, S., Liu, L., & Kaplan, H. J. (2010). Epithelial-mesenchymal transition and proliferation of retinal pigment epithelial cells initiated upon loss of cell-cell contact. Invest Ophthalmol Vis Sci, 51(5), 2755–2763. https://doi.org/10.1167/iovs.09-4725

Tangeman, J. A., Luz-Madrigal, A., Sreeskandarajan, S., Grajales-Esquivel, E., Liu, L., Liang, C., Tsonis, P. A., & Del Rio-Tsonis, K. (2021). Transcriptome Profiling of Embryonic Retinal Pigment Epithelium Reprogramming. Genes (Basel), 12(6). https://doi.org/10.3390/genes12060840

Tazaki, A., Tanaka, E. M., & Fei, J. F. (2017). Salamander spinal cord regeneration: The ultimate positive control in vertebrate spinal cord regeneration. Dev Biol, 432(1), 63–71. https://doi.org/10.1016/j.ydbio.2017.09.034

Team, R. (2020). RStudio: Integrated Development for R. http://www.rstudio.com/.

Thummel, R., Kassen, S. C., Enright, J. M., Nelson, C. M., Montgomery, J. E., & Hyde, D. R. (2008). Characterization of Muller glia and neuronal progenitors during adult zebrafish retinal regeneration. Exp Eye Res, 87(5), 433–444. https://doi.org/10.1016/j.exer.2008.07.009

Tsonis, P. A., & Del Rio-Tsonis, K. (2004). Lens and retina regeneration: transdifferentiation, stem cells and clinical applications. Exp Eye Res, 78(2), 161–172. https://doi.org/10.1016/j.exer.2003.10.022

Tucker, B., Klassen, H., Yang, L., Chen, D. F., & Young, M. J. (2008). Elevated MMP Expression in the MRL Mouse Retina Creates a Permissive Environment for Retinal Regeneration. Invest Ophthalmol Vis Sci, 49(4), 1686–1695. https://doi.org/10.1167/iovs.07-1058

UniProt, C. (2021). UniProt: the universal protein knowledgebase in 2021. Nucleic Acids Res, 49(D1), D480–D489. https://doi.org/10.1093/nar/gkaa1100

Vergara, M. N., & Del Rio-Tsonis, K. (2009). Retinal regeneration in the Xenopus laevis tadpole: a new model system. Mol Vis, 15, 1000–1013. https://www.ncbi.nlm.nih.gov/pubmed/19461929

Walcott, J. C., & Provis, J. M. (2003). Muller cells express the neuronal progenitor cell marker nestin in both differentiated and undifferentiated human foetal retina. Clin Exp Ophthalmol, 31(3), 246–249. https://doi.org/10.1046/j.1442-9071.2003.00638.x

Wan, J., & Goldman, D. (2016). Retina regeneration in zebrafish. Curr Opin Genet Dev, 40, 41–47. https://doi.org/10.1016/j.gde.2016.05.009

Xia, X., Teotia, P., & Ahmad, I. (2018). Lin28a regulates neurogliogenesis in mammalian retina through the Igf signaling. Dev Biol, 440(2), 113–128. https://doi.org/10.1016/j.ydbio.2018.05.007

Yoshii, C., Ueda, Y., Okamoto, M., & Araki, M. (2007). Neural retinal regeneration in the anuran amphibian Xenopus laevis post-metamorphosis: transdifferentiation of retinal pigmented epithelium regenerates the neural retina. Dev Biol, 303(1), 45–56. https://doi.org/10.1016/j.ydbio.2006.11.024

Young, M. D., Wakefield, M. J., Smyth, G. K., & Oshlack, A. (2010). Gene ontology analysis for RNA-seq: accounting for selection bias. Genome Biol, 11(2), R14. https://doi.org/10.1186/gb-2010-112-r14

Zhao, S., Rizzolo, L. J., & Barnstable, C. J. (1997). Differentiation and transdifferentiation of the retinal pigment epithelium. Int Rev Cytol, 171, 225–266. https://doi.org/10.1016/s0074-7696(08)62589-9

Zhou, M., Geathers, J. S., Grillo, S. L., Weber, S. R., Wang, W., Zhao, Y., & Sundstrom, J. M. (2020). Role of Epithelial-Mesenchymal Transition in Retinal Pigment Epithelium Dysfunction. Front Cell Dev Biol, 8, 501. https://doi.org/10.3389/fcell.2020.00501

